# Reconstructing B cell receptor sequences from short-read single cell RNA-sequencir with BRAPeS

**DOI:** 10.1101/389999

**Authors:** Shaked Afik, Gabriel Raulet, Nir Yosef

**Affiliations:** Center for Computational Biology, University of California, Berkeley, Berkeley, CA, 94720, USA; Department of Computer Science, University of California, Davis, Davis, CA, 95616, USA; Department of Electrical Engineering and Computer Science, University of California, Berkeley, Berkeley, CA, 94720, USA; Ragon Institute of MGH, MIT and Harvard, Cambridge, MA, USA; Chan Zuckerberg Biohub, San Francisco, CA 94158, USA

## Abstract

RNA-sequencing of single B cells provides simultaneous measurements of the cell state and its binding specificity. However, in order to uncover the latter further reconstruction of the B cell receptor (BCR) sequence is needed. We present BRAPeS, an algorithm for reconstructing BCRs from short-read paired-end single cell RNA-sequencing. BRAPeS is accurate and achieves a high success rate even at very short (25bp) read length, which can decrease the cost and increase the number of cells that can be analyzed compared to long reads. BRAPeS is publicly available in the following link: https://github.com/YosefLab/BRAPeS.

## INTRODUCTION

B cells play a significant role in the adaptive immune system, providing protection against a wide range of pathogens. This diversity is due to the B cell receptor (BCR), which enables different cells to bind different pathogens (Imkeller & Wardemann, 2018). Single cell RNA-sequencing (scRNA-seq) has emerged as one of the leading technologies to characterize and study heterogeneity in the immune system across cell types, development and dynamic processes (Papalexi & Satija, 2018; Villani *et al*, 2018). Combining transcriptome analysis with BCR reconstruction in single cells can provide valuable insights to the relation between BCR and cell state, as was demonstrated by similar studies in T cells (Afik *et al*, 2017; Stubbington *et al*, 2016; Eltahla *et al*, 2016).

The BCR is comprised of two chains, a heavy chain (IgH) and a light chain (IgL, either a kappa or a lambda chain). Each chain is encoded in the germline by multiple segments of three types - variable (V), joining (J) and constant (C) segments (the heavy chain also includes a diversity (D) segment, see Materials and Methods). The specificity of the BCRs comes from the V(D)J recombination process, in which for each chain one variable (V) and one joining (J) segments are recombined in a process which introduces mutations, insertions and deletions into the junction region between the segments, called the complementarity determining region 3 (CDR3) (Tonegawa, 1983). The resulting sequence is the main determinant of the cell’s ability to recognize a specific antigen. Following B cell activation, somatic hypermutations are introduced in the complementarity determining regions of the BCR, and the constant region is replaced in a process termed isotype switching or class switching (Di Noia & Neuberger, 2007). The random mutations make BCR reconstruction a challenging task. While methods to reconstruct BCR sequences from full length scRNA-seq are available (Canzar *et al*, 2017; Rizzetto *et al*, 2018; Lindeman *et al*, 2018) (as well as single cell V(D)J enriched libraries from 10x Genomics (Single Cell Immune Profiling - 10x Genomics)), they were only tested on long reads (150bp and 50bp). The ability to reconstruct BCR sequences from short length (25-30bp) reads is important, as it can decrease cost which can, in turn, increase the number of cells which could be feasibly analyzed.

We introduce BRAPeS (“BCR Reconstruction Algorithm for Paired-end Single cells”), an algorithm and software for BCR reconstruction. Conversely to other methods, BRAPeS was designed to work with short (25-30bp) reads, and indeed we demonstrate that under these settings it performs better than other methods. Furthermore, we show that the performance of BRAPeS when provided with short reads is similar to what can be achieved with much longer (50-150bp) reads from the same cells, suggesting that BCR reconstruction does not necessitate costly sequencing with many cycles.

## RESULTS

BRAPes is an extension of the TCR reconstruction software TRAPeS (Afik *et al*, 2017), with significant modifications added to address the processes of class switching and somatic hypermutations which are specific to B cells (Figure 1, see Materials and Methods for full description of BRAPeS). Briefly, BRAPeS takes as input the alignment of the reads to the genome. BRAPeS first recognizes the possible V and J segments by finding reads with one mate mapping to a V segment and the other mate mapping to a J segment. BRAPeS then collects all unmapped reads whose mates were mapped to the V/J/C segments, assuming that most CDR3-originating reads will be unmapped when aligning to the reference genome. Then, BRAPeS reconstructs the CDR3 region with an iterative dynamic programming algorithm. At each step, BRAPeS aligns the unmapped reads to the edges of the V and J segments, using the sequence of the aligned reads to extend the V and J sequences until convergence. Next, BRAPeS determines the BCR isotype by appending all possible constant segments to the reconstructed sequence and taking the most likely complete transcript based on transcriptomic alignment with RSEM (Li & Dewey, 2011). Finally, BRAPeS corrects for somatic hypermutations by collecting all reads aligning to the genomic CDR1 and CDR2 sequences and aligning the reads against themselves to obtain a reconstruction of the consensus sequence. BRAPeS determines the CDR3 sequences of all BCRs and its productivity based on the criteria established by the international ImMunoGeneTic information system (IMGT) (Lefranc *et al*, 2015; Lefranc, 2014). BRAPeS reports the full BCR sequences per cell, their productivity status, V/J/C segments and the number of reads mapped to the various segments of the BCRs.

**Figure 1:**
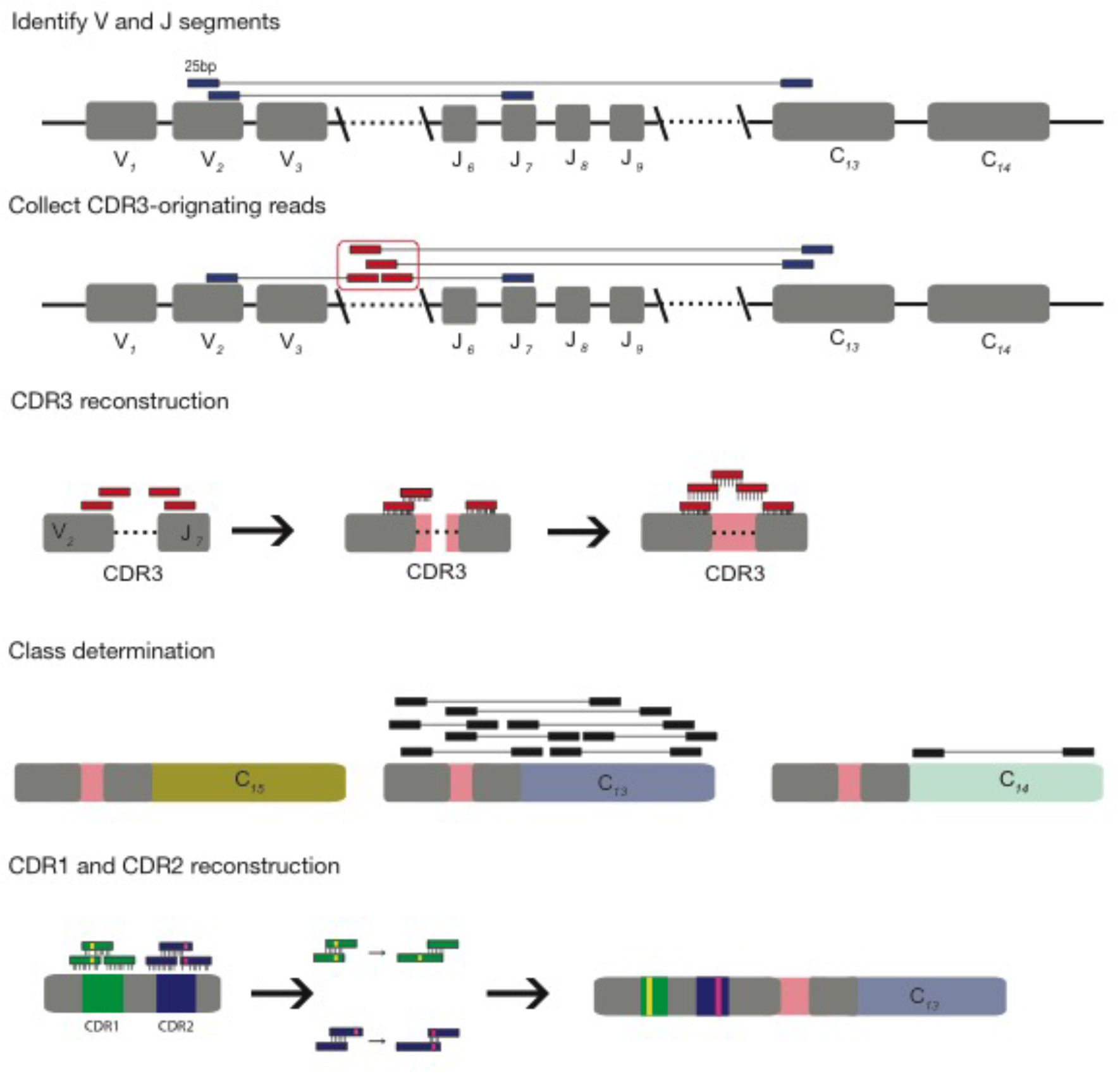
The BRAPeS algorithm. First, the V and J segments are selected by searching for paired reads with one read mapping to a V segment and its mate mapping to a J segment. Next, putative CDR3-originating reads are identified as the unmapped reads whose mates map to the V/J/C segments. BRAPeS runs an iterative dynamic programming algorithm to align the CDR3-originating reads to the V and J segments and extend them until they overlap. BRAPeS then determines the BCR isotype by running RSEM on all possible full BCR transcripts (the reconstructed V-J segments combined with all possible constant segments). Finally, BRAPeS corrects the CDR1 and CDR2 regions for somatic hypermutations by building a consensus sequence of the reads aligning to these regions.

We evaluated BRAPeS’ performance on 374 cells from two previously published data sets - 174 human B cells and 200 mouse B cells (Materials and Methods, Additional File 1) (Canzar *et al*, 2017; Wu *et al*, 2016). To evaluate BRAPes, we first trimmed the original reads (50bp for the human data and 150bp for the mouse data) and kept only the outer 25 or 30 bases. We compared BRAPeS’ performance on the trimmed data to two other previously published methods - BASIC (Canzar *et al*, 2017) and VDJPuzzle (Rizzetto *et al*, 2018) applied either on the trimmed data or the original long reads.

When applied to 30bp reads, BRAPeS’ success rates are similar to other methods for the light chain but are higher for heavy chain reconstruction (Figure 2a, Table S1). BRAPeS reconstructs productive heavy chains in a total of 349 cells, 93.3% of the cells across both datasets, and reconstructs productive light chains in 364 cells (97.3% of the cells). These results are in line with the success rates of BASIC and VDJPuzzle on the original long reads: BASIC reconstructs productive heavy and light chains in 352 (94.1%) and 364 (97.3%) cells, respectively, and VDJPuzzle reconstructs heavy chains in 346 (92.5%) cells and light chains in 368 (98.4%) cells. On 30bp reads, BASIC and VDJPuzzle achieve reconstruction rates similar and even slightly higher compared to long reads for the light chain (362 (96.8%) cells and 370 (98.9%) cells with a productive light chain in BASIC and VDJPuzzle, respectively). However, BASIC and VDJPuzzle see a decline in success rates for the heavy chain, reconstructing a productive heavy chain in only 273 (73%) cells for BASIC and 242 (64.7%) cells for VDJPuzzle (Figure 2a, Table S1).

**Figure 2:**
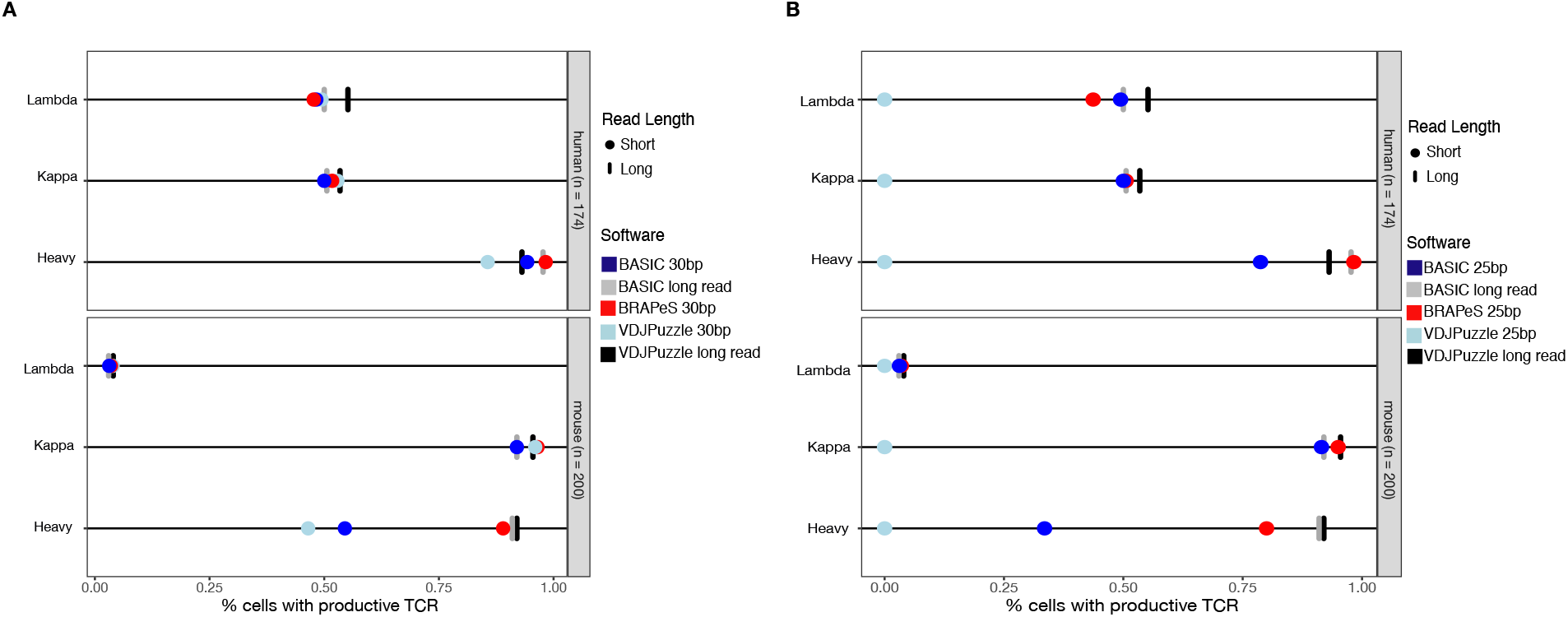
BRAPeS success rates. **A)** Fraction of cells with a successful reconstruction of a productive CDR3 in human and mouse B cells using the following methods: VDJPuzzle applied to the original, long read data (black line) and the trimmed version of the data, trimmed to 30bp (light blue circle). BASIC applied to the long read (grey line) and the trimmed data (dark blue circle), and BRAPeS applied to the trimmed data (red circle). **B)** Same as A, but the trimmed version of the data was trimmed down to include only the outer 25bp, instead of 30bp.

BRAPeS is also able to maintain a high success rate on 25bp reads, reconstructing heavy chains in 331 (88.5%) cells and light chains in 357 (95.5%) cells (Figure 2b and Table S2). Yet, we observe a substantial decrease in the results of other methods. VDJPuzzle is unable to reconstruct any chains with 25bp reads. This is likely due to its use of the *De-novo* assembler Trinity (Grabherr *et al*, 2011) which requires a seed k-mer length of 25bp that is unsuitable for very short reads. Similarly to 30bp, BASIC is able to maintain a high reconstruction rate for light chains, with productive reconstructions in 363 (97.1%) cells, but is only able to reconstruct productive heavy chains in 204 (54.6%) cells (Figure 2b, Table S2). Moreover, BASIC only outputs fasta sequences, thus requiring further processing to annotate the BCR.

We next turn to evaluate the accuracy of the short-read based CDR3 reconstructions, by comparing the resulting sequences to those obtained with long reads (Figure 3, Materials and Methods). We use the long-read based reconstruction of BASIC as a reference (we achieve similar results with VDJPuzzle on the long read data; Figure S1) and evaluate the accuracy in terms of sensitivity (how many of the CDR3 sequences in the full length data have an identical reconstruction with the short-reads) and specificity (how many of the CDR3 sequences in the short-read data have an identical long-read reconstruction). In general, all methods show a high level of specificity, having almost all CDR3 sequences identical to the sequences reconstructed on long reads, whenever both read lengths produce a productive reconstruction (Figure 3a-b). In accordance with the higher success rate, BRAPeS shows a high sensitivity, with an average rate of 0.96 for 30bp data and 0.93 for 25bp data (Figure 3c-d). This is in line with the agreement of different methods on the original data, as VDJPuzzle on long reads has an average sensitivity rate of 0.96. On the trimmed data, BASIC and VDJPuzzle show a lower sensitivity rate - BASIC achieves sensitivity rates of 0.91 and 0.85 for 30bp and 25bp respectively, and VDJPuzzle has a sensitivity rate of 0.89 with the 30bp data.

**Figure 3:**
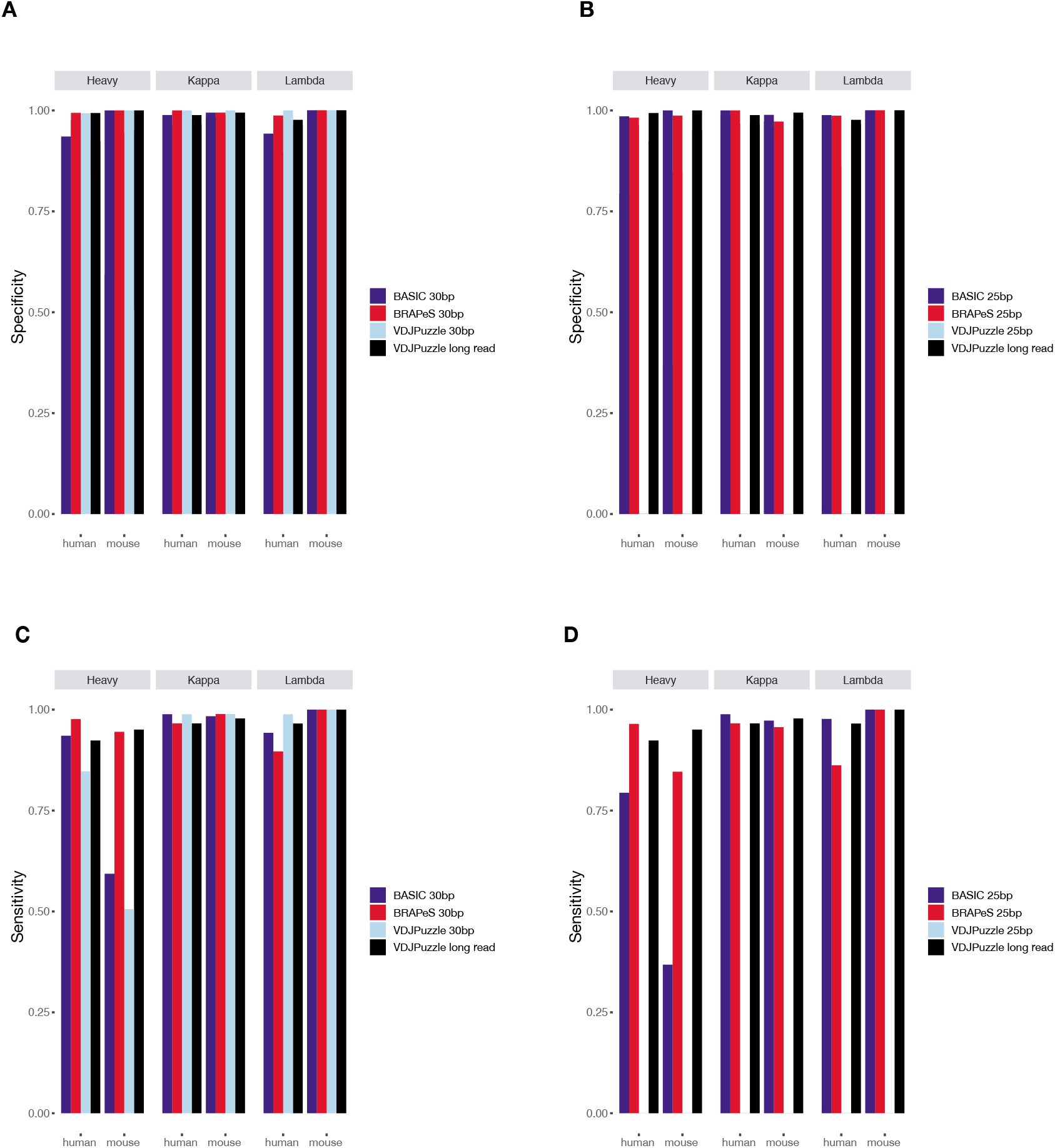
Sensitivity and specificity of BRAPeS. **A)** Specificity of BRAPeS for 30bp. The fraction of cells with a CDR3 sequence identical to the CDR3 reconstructed by BASIC on the long-read data, using the following methods: VDJPuzzle when applied to the long-read data (black), BRAPeS (red), BASIC (dark blue) and VDJPuzzle (light blue) applied to a version of the data trimmed to 30bp. The fraction is calculated only for cells that had a productive chain in both the long read BASIC results and the other method. **B)** Specificity of BRAPeS for 25bp. Same as A, except the short-read version of the data was trimmed to include only the outer 25bp, instead of 30bp. **C)** Sensitivity of BRAPeS for 30bp. Same as A, except the fraction is calculated out of all the cells that had a productive chain when running BASIC on the long-read data. **D)** Sensitivity of BRAPeS for 25bp. Same as B, except the fraction is calculated out of all the cells that had a productive chain when running BASIC on the long-read data.

Coupling BCR reconstruction with transcriptome analysis in single cells can provide valuable information about the effect of binding specificity and isotype to cellular heterogeneity. BRAPeS is a software for BCR reconstruction which utilizes short read scRNA-seq, allowing for decreased cost. BRAPeS is accurate and has a higher success rate on short reads compared to existing methods, especially for 25bp reads and heavy chains. BRAPeS is publicly available at https://github.com/YosefLab/BRAPeS

## MATERIALS AND METHODS

### The BRAPeS algorithm

The input given to BRAPeS is a directory where each subdirectory includes genomic alignments of a single cell.

The BRAPeS algorithm have several steps, performed separately for each chain in each cell:

1. **Identifying possible pairs of V and J segments:** BRAPeS searches for reads where one mate of the pair is mapped to a V segment and the other mate is mapped to a J segment. BRAPeS collects all possible V-J pairs and will try to reconstruct complete BCRs from all possible pairs. Since the D segment is very short, reads do not align to it, thus as part of the reconstruction step (step 3) BRAPeS will also reconstruct the sequence of the D segment. If no V-J pairs are found, BRAPeS will look for V-C and J-C pairs and will take all possible V/J pairing of the found V and J segments. In case of many possible V-J pairs (which can occur due to the similarity among the segments), the user can limit the number of V-J pairs to attempt reconstruction on. BRAPeS will rank the V-J pairs based on the number of reads mapped to them and take only the top few pairs (the exact number is a parameter controlled by the user).
2. **Collecting the set of putative CDR3-originating reads:** BRAPeS collects the set of reads that are likely to originate from the CDR3 region. Those are the reads that are unmapped to the reference genome, but their mates are mapped to the V/J/C segments. In addition, since the first step of CDR3 reconstruction includes alignment to the ends of the genomic V and J sequences, BRAPeS also collects the reads mapping to the V and J segments.
3. **Reconstructing the CDR3 region:** For each V-J pair, BRAPeS extends the edges of the V and J segments with an iterative dynamic programming algorithm. BRAPeS starts the reconstruction from the end bases of the V and J segment (3’ end of the V segment and 5’ end of the J segment). The number of bases is a parameter which can be controlled by the user, set by default to the length of the J segment. In each iteration, BRAPeS tries to align all the unmapped reads to the V and J segments separately with the Needleman-Wunsch algorithm with the following scoring scheme: +1 for match, −1 for mismatch, −20 for gap opening and −4 for gap extension. In addition, BRAPeS does not penalize having a read “flank” the genomic segment. All reads that passed a user defined threshold are considered successful alignments. BRAPeS will then build the extended V and J segments by taking for each position the base which appears in most reads. BRAPeS will continue to run this process for a given number of iterations or until the V and J segments overlap. BRAPeS can also run a “one-side” mode, where if an overlap was not found (e.g. due to assigning the wrong V segment), BRAPeS will attempt to determine the productivity of only the extended V and only of the extended J segment.
4. **Isotype determination:** To find the BCR class, for each V-J pair with a reconstructed CDR3, BRAPeS concatenates all possible constant segments. Then, BRAPeS runs RSEM (Li and Dewey, 2011) on all sequences using all the paired-end reads with at least one mate that was mapped the genomic V/J/C segments as input. For each V-J pair BRAPeS takes the constant region with the highest expected count as the chosen constant segment.
5. **CDR1 and CDR2 reconstruction:** All the reads from step 4 are aligned against the genomic CDR1 and CDR2 sequences obtained from IMGT using the C++ SeqAn package (Döring *et al*, 2008). The genomic CDR1 and CDR2 sequences are optionally extended into the adjacent framework regions based on an input parameter (in order for reads that flank the CDR1 or CDR2 to be successfully identified). Reads scoring above a user defined threshold in a global alignment (using the same scoring scheme as step 3) are saved for reconstruction. The reconstruction runs ten randomized iterations of the following algorithm: reads are randomly selected for global alignment against each other. Alignments whose matching segments are perfect alignments recursively extend the reconstructed sequence. Once all reads have been aligned or discarded, the reconstructed sequence is aligned against the genomic sequence to determine its starting and ending points. The most frequent reconstruction is selected as the final reconstruction for the CDR1 and CDR2.
6. **Separating similar BCRs and determining chain productivity:** After selecting the top isotype for each V-J pair, BRAPeS determines if the reconstructed sequence is productive (i.e. the V and J are in the same reading frame with no stop codon in the CDR3) and annotates the CDR3 junction. If more than one V-J pair produces a CDR3 sequence (either due to having more than one recombined chain in the cell or due to similar V-J segments resulting in the same CDR3 sequence reconstruction), BRAPeS will rank the various reconstructions based on their expression values from RSEM.

The output for BRAPeS is the full ranked list of reconstructed chains, including the CDR3 sequences, V/J/C annotations and the number of reads mapped to each segment, as well as a summary file of the success rates across all cells.

BRAPeS is implemented in python. To increase performance, the dynamic programming algorithm is implemented in C++ using the SeqAn package (Döring et al., 2008). Moreover, to decrease running time for deeply sequenced cells, BRAPeS has the option to randomly downsample the V-J aligning reads and the putative CDR3-originating reads to only 10,000 reads. BRAPeS is publicly available and can be downloaded in the following link: https://github.com/YosefLab/BRAPeS

### Data preprocessing

Raw fastq files of mouse B cells were downloaded from Wu et al. (ArrayExpress E-MTAB-4825) (Wu et al., 2016). All analysis was performed on the 200 cells that were available through ArrayExpress. Raw fastq files for the human data from Canzar et al. (Canzar et al., 2017) were provided by the author. We excluded single-end cells and cells filtered out in the original study, leaving a total of 174 cells. Next, reads were trimmed to be 25 or 30bp paired-end with trimmomatic (Bolger et al., 2014), keeping only the outer bases.

For BRAPeS, low quality reads were trimmed using trimmomatic with the following parameters: LEADING:15, TRAILING:15, SLIDINGWINDOW:4:15, MINLEN:16. The remaining reads were aligned to the genome (hg38 or mm10) using Tophat2 (Kim et al., 2013).

### Running BRAPeS

For this study, BRAPeS was run using the following parameters for the human data: “-score 15 - top 6 -byExp -iterations 6 -downsample -oneSide -HVR_extension 15 -HVR_score 15 - HVR_moveOn 30000 -HVR_max_reads 200”, and with the following parameters for the mouse data: “-score 15 -oneSide -byExp -top 10 -HVR_extension 15 -HVR_score 15 -HVR_moveOn 30000 -HVR_max_reads 200”. In addition, as some cells required a higher alignment score threshold, we ran BRAPeS with a scoring threshold of 21 for cells without a productive chain.

### Running VDJPuzzle and BASIC

We ran VDJPuzzle using default parameters, providing VDJPuzzle with the hg38 genome and GRCh38.p2 annotation for human, and mm10 genome with the GRCm38.p4 annotation for mouse. We then considered only results which appeared in the “summary_corrected” folder as valid productive reconstructions.

BASIC was ran with default parameters. After running BASIC we collected all the output fasta files and ran them through IMGT/HighV-Quest (Alamyar et al., 2012; Li et al., 2013). Only sequences that resulted in productive CDR3 according to IMGT were considered successful reconstructions.

### Comparison of sensitivity and specificity

To determine the accuracy of the methods, we compared the reconstructed CDR3 amino acid sequences to the reconstruction produced by running BASIC or VDJPuzzle on the long reads. Only CDR3s with amino acid sequences identical to the sequences reconstructed on the long-read data were considered accurate. In case of more than one reconstructed CDR3 sequence, if both methods had at least one identical CDR3 sequence it was considered an accurate reconstruction.

## ACKNOWLEDGEMENTS

This work was supported by US National Institute of Health, grant number 5U19AI090023-07

## AUTHOR CONTRIBUTIONS

S.A. wrote BRAPeS, performed the analysis and wrote the manuscript, G.R. designed and wrote the code for CDR1 and CDR2 reconstruction. N.Y. designed and oversaw the study and wrote the manuscript.

## DATA AVAILABILITY

Mouse raw fastq files from Wu et al. were downloaded from ArrayExpress (E-MTAB-4825). Human raw fastq files from Canzar et al. were provided by the author. BRAPeS can be downloaded in the following link: https://github.com/YosefLab/BRAPeS

## ADDITIONAL FILES

**Additional file 1:** BRAPeS output for datasets analyzed in this study. In addition to the standard output, the last column mentions the alignment threshold used for each reconstruction

---

**Figure S1:**
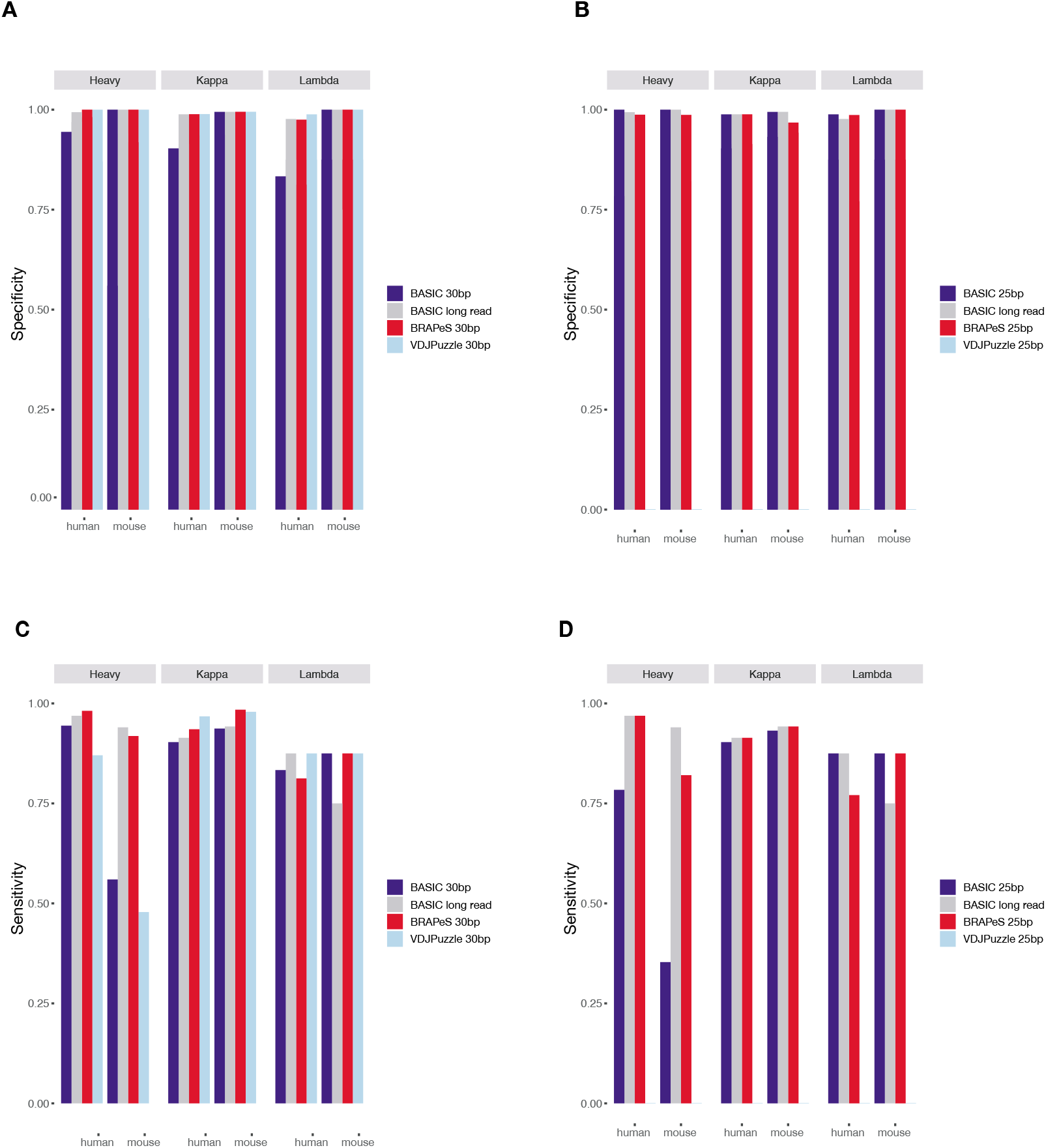
Sensitivity and specificity of BRAPeS compared to VDJPuzzle reconstructions on long read data. **A)** Specificity of BRAPeS for 30bp. The fraction of cells with a CDR3 sequence identical to the CDR3 reconstructed by VDJPuzzle on the long read data, using the following methods: BASIC when applied to the long read data (grey), BRAPeS (red), BASIC (dark blue) and VDJPuzzle (light blue) applied to a version of the data trimmed to 30bp. The fraction is calculated only for cells that had a productive chain in both the long read VDJPuzzle results and the other method. **B)** Specificity of BRAPeS for 25bp. Same as A, except the short read version of the data was trimmed to include only the outer 25bp, instead of 30bp. **C)** Sensitivity of BRAPeS for 30bp. Same as A, except the fraction is calculated out of all the cells that had a productive chain when running VDJPuzzle on the long read data. **D)** Sensitivity of BRAPeS for 25bp. Same as B, except the fraction is calculated out of all the cells that had a productive chain when running VDJPuzzle on the long read data.

**Table S1:** Detailed description of the number of productive reconstructions for the original long reads and 30bp sequencing

|  | **Human (n = 174)** |  |  | **Mouse (n = 200)** |  |  |
| --- | --- | --- | --- | --- | --- | --- |
|  | **Heavy** | **Light**<br>**(kappa or**<br>**Lambda)** | **Both**<br>**kappa and**<br>**Lambda** | **Heavy** | **Light**<br>**(kappa or**<br>**Lambda)** | **Both kappa**<br>**and**<br>**Lambda** |
| BASIC - long read | 170 (97.7%) | 174 (100%) | 1 (0.6%) | 182 (91%) | 190 (95%) | 0 (0%) |
| VDJPuzzle - long read | 162 (93.1%) | 172 (98.9%) | 17 (9.77%) | 184 (92%) | 196 (98%) | 3 (1.5%) |
| BRAPeS - 30bp | 171 (98.3%) | 165 (94.8%) | 8 (4.6%) | 178 (89%) | 199 (99.5%) | 1 (0.5%) |
| BASIC - 30bp | 164 (94.3%) | 171 (98.3%) | 0 (0%) | 109 (54.5%) | 191 (95.5%) | 0 (0%) |
| VDJPuzzle - 30bp | 149 (85.6%) | 172 (98.9%) | 6 (3.45%) | 93 (46.5%) | 198 (99%) | 1 (0.5%) |

**Table S2:** Detailed description of the number of productive reconstructions for the original long reads and 25bp sequencing

|  | Human (n = 174) |  |  | Mouse (n = 200) |  |  |
| --- | --- | --- | --- | --- | --- | --- |
|  | Heavy | Light<br>(kappa or<br>Lambda) | Both<br>kappa and<br>Lambda | Heavy | Light<br>(kappa or<br>Lambda) | Both kappa<br>and<br>Lambda |
| BASIC - long read | 170 (97.7%) | 174 (100%) | 1 (0.6%) | 182 (91%) | 190 (95%) | 0 (0%) |
| VDJPuzzle - long read | 162 (93.1%) | 172 (98.9%) | 17 (9.77%) | 184 (92%) | 196 (98%) | 3 (1.5%) |
| BRAPeS - 25bp | 171 (98.3%) | 161 (92.5%) | 3 (1.72%) | 160 (80%) | 196 (98%) | 1 (0.5%) |
| BASIC - 25bp | 137 (78.7%) | 173 (99.4%) | 0 (0%) | 67 (33.5%) | 190 (95%) | 0 (0%) |
| VDJPuzzle -25bp | 0 (0%) | 0 (0%) | 0 (0%) | 0 (0%) | 0 (0%) | 0 (0%) |

